# Eye drift during fixation predicts visual acuity

**DOI:** 10.1101/2022.03.21.485131

**Authors:** Ashley M. Clark, Janis Intoy, Michele Rucci, Martina Poletti

## Abstract

Visual acuity is commonly assumed to be determined by the eye optics and spatial sampling in the retina. Unlike a camera, however, the eyes are never stationary during the acquisition of visual information, a jittery motion known as ocular drift, incessantly displaces stimuli over many photoreceptors. Previous studies have shown that acuity is impaired in the absence of retinal image motion caused by eye drift. However, the relation between individual drift characteristics and acuity remains unknown. Here we show that (*a*) healthy emmetropes exhibit a large variability in their amount of drift; and (*b*) that these differences profoundly affect the structure of spatiotemporal signals to the retina. We further show that (*c*) the spectral distribution of the resulting luminance modulations strongly correlates with individual visual acuity; and (*d*) that natural inter-trial fluctuations in the amount of drift modulate acuity. As a consequence, in healthy emmetropes acuity can be predicted from the motor behavior elicited by a simple fixation task, without directly measuring it. These results shed new light on how oculomotor behavior contributes to fine spatial vision.

**Significance:** Healthy humans can visually resolve extremely fine patterns, in some cases with the relevant features spanning less than a single photoreceptor on the retina. This accomplishment is particularly remarkable considering that the eyes are never stationary. Ocular drift—a motion that eludes human awareness—shifts the stimulus across many photoreceptors during the acquisition of visual information. Here we show that visual acuity depends on ocular drift. Natural variations in the amount of drift are associated with acuity both within and across subjects, so that individual acuity limits can be directly inferred from the amount of motion during fixation on a marker. Results closely follow the strength of the luminance modulations caused by ocular drift, providing support to long-standing dynamic theories of visual acuity.

## Introduction

Fine spatial vision is important for normal functioning. When vision in the central fovea—the region where photoreceptors are most densely packed—is compromised, many daily activities are severely impacted^1,2^. It is therefore not surprising that visual acuity, the spatial resolving capacity of the visual system ^3^, is assessed in *nearly every* eye exam. Although acuity varies considerably among healthy individuals, emmetropic observers are typically capable of discriminating patterns at frequencies of ∼30 cycles/deg and some can resolve up to 60 cycle/deg. This capability is remarkable considering that the eye is never stationary during the acquisition of visual information. In the interval in between saccades, a persistent jittery motion, known as ocular drift, incessantly displace the stimulus on the retina across many photoreceptors, even when looking at a single point ^4–6^ (Fig. 1*A*-*B*). This motion raises fundamental questions on the mechanisms underlying the establishment of fine spatial representations and how the visual system avoids the perceptual blur that one would expect from the smearing of the stimulus on the retina ^7–10^. It challenges the traditional, purely spatial view that regards acuity limits as only determined by optical and anatomical factors ^11–15^.

**Figure 1:**
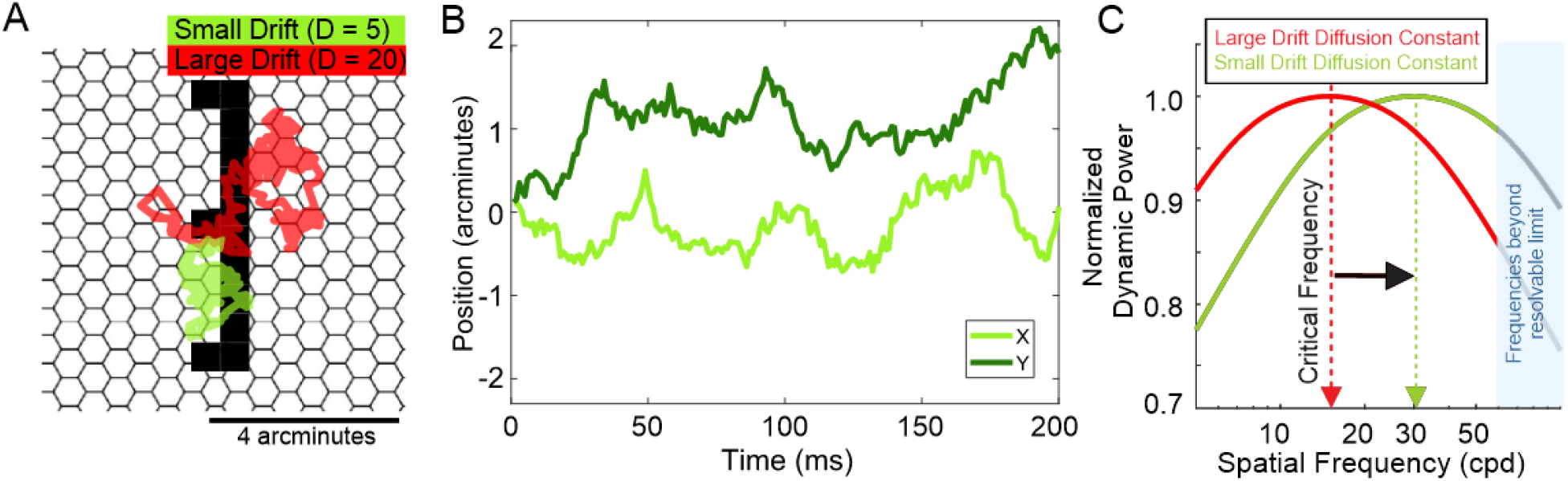
Ocular Drift. (**A-B**) Examples of typical ocular drifts exhibited by two observers while examining the stimuli of this study. These subjects differ in the extent of their drifts (green and red traces). In **A**, traces are superimposed to the stimulus and shown relative to a schematic of the photoreceptor mosaic in the central fovea drawn to scale. The drift diffusion constants are reported on top. In **B**, the vertical and horizontal components of the green trace in *A* are shown as a function of time. (**C**) Average amplitudes of the luminance fluctuations resulting from these two types of drift during viewing of gratings at individual spatial frequencies. Each curve represents the signal effective in driving retinal ganglion cells, estimated as the total power released by drift at non-zero temporal frequencies within the bandwidth of neuronal sensitivity. Note that, rather than being constant across spatial frequencies, this signal is strongest within a frequency band that varies with the amount of drift ^33,28^. The shaded blue region marks frequencies beyond human sensitivity ^34^.

An alternative view has long been proposed. It has been argued for over a century that the fixational motion of the eye may actually be beneficial to visual acuity by *structuring*—rather than just *refreshing*—neural activity ^16–21^, an hypothesis originally formulated by Hering ^16^ and later refined into the so-called dynamic theories of visual acuity ^18,19,17^. This idea that both the temporal and spatial component of the visual input need to be taken into account is well established in models explaining motion sensitivity ^22^, yet it has encountered resistance when applied to spatial perception. In fact, early behavioral experiments did not support these proposals ^23–25^. However, these pioneering experiments also suffered from significant technological and methodological limitations (see Rucci et al, 2007 for a discussion of issues). In contrast, recent studies that used more advanced and flexible methods for controlling retinal stimulation have shown that fine spatial vision is specifically impaired in the absence of the retinal image motion resulting from ocular drift ^26–28^. Furthermore, it has been observed that the way drift transforms spatial patterns into a spatiotemporal flow on the retina implements a crucial information-processing step tuned to the characteristics of the natural visual world ^6^. This transformation discards redundant information and enhances neural responses to luminance discontinuities, processes long argued to be important goals of early visual processing ^29,25,30,21,31^, which is expected given that neurons in the retina and the early visual system are relatively insensitive to an unchanging input ^32^.

The spatio-temporal reformatting of visual input signals resulting from ocular drift carries important implications for visual acuity. It is now known that, as the eye drifts, the range of spatial frequencies in which the resulting luminance modulations possess maximum strength depends on the extent of ocular motion ^33^, with smaller drifts (slower and more curved motion) shifting the power of input signals toward progressively higher ranges of spatial frequencies (Fig. 1*C*). The recent observation that the amount of drift is reduced in a standard acuity test ^28^ suggests that humans actively attempt to take advantage of this input reformatting. However, it remains unclear how ocular drift interacts with individual constraints imposed by the eye optics and retinal anatomy. It is known that, like visual acuity, the extent of ocular drift also varies considerably and idiosyncratically across healthy emmetropes ^5^. Yet, no previous study has examined whether a relation exists between these two variables and whether acuity is modulated by the specific amount of ocular drift.

Since the amount of ocular drift regulates the spatial frequency bandwidth with strongest luminance modulations, use of this signal yields specific predictions on how eye motion and acuity limits interact both within and across subjects. Specifically, to take maximal advantage of drift-induced modulations, one would expect emmetropic observers to tune their drifts according to individual acuity limits, so that the smallest drifts are exhibited by subjects with the highest acuity. Furthermore, in individual observers, acuity is expected to vary according to the patterns of drift exhibited at any given time. Here we test these predictions. Ocular drift is typically neglected or not assessed in experimental and clinical examinations of visual acuity. Confirmation of these hypotheses will emphasize the need to pay attention to eye movements at this scale and provide further support to the notion that fine spatial vision is an active process that relies on oculomotor-induced temporal modulations.

## Results

To quantify predictions on how eye movements may influence acuity, we first examined the luminance signals resulting from ocular drift. To this end, we assume drift to be well approximated by a Brownian Motion (BM) process. This model of drift has been used extensively in the literature ^35,36,9^, and a large body of previous findings indicates that the BM approximation is well justified ^35,33,28^. This model is also convenient as it summarizes the contributions of both velocity and curvature of eye movements into a single parameter, the diffusion constant (*D*), which describes how rapidly the line of sight moves away from its current location.

Fig. 2 describes how changes in the stimulus and in the amount of drift affect visual input signals. As the eye drifts, the external spatial stimulus is converted into a spatio-temporal luminance flow on the retina, the characteristics of which depend on both the stimulus and the extent of drift. The top row of Fig. 2*A* provides examples of the luminance modulations resulting with exposure to stimuli at two different spatial frequencies, 10 and 30 cycles/deg, while drifting by a small amount (*D* = 5 arcmin^2^/s). As the spatial frequency of the stimulus increases, modulations increase both in amplitude and temporal frequency bandwidth—*i*.*e*., how rapidly they change. Enlarging drift has a similar effect to increasing the spatial frequency of the stimulus. With a drift magnified by four times (*D* = 20 arcmin^2^/s), larger and faster luminance modulations are visible with the 10 cycles/deg stimulus. With the 30 cycles/deg stimulus, these fluctuations are so rapid that many are likely beyond the temporal sensitivity of retinal neurons (bottom row of Fig. 2*A*).

**Figure 2:**
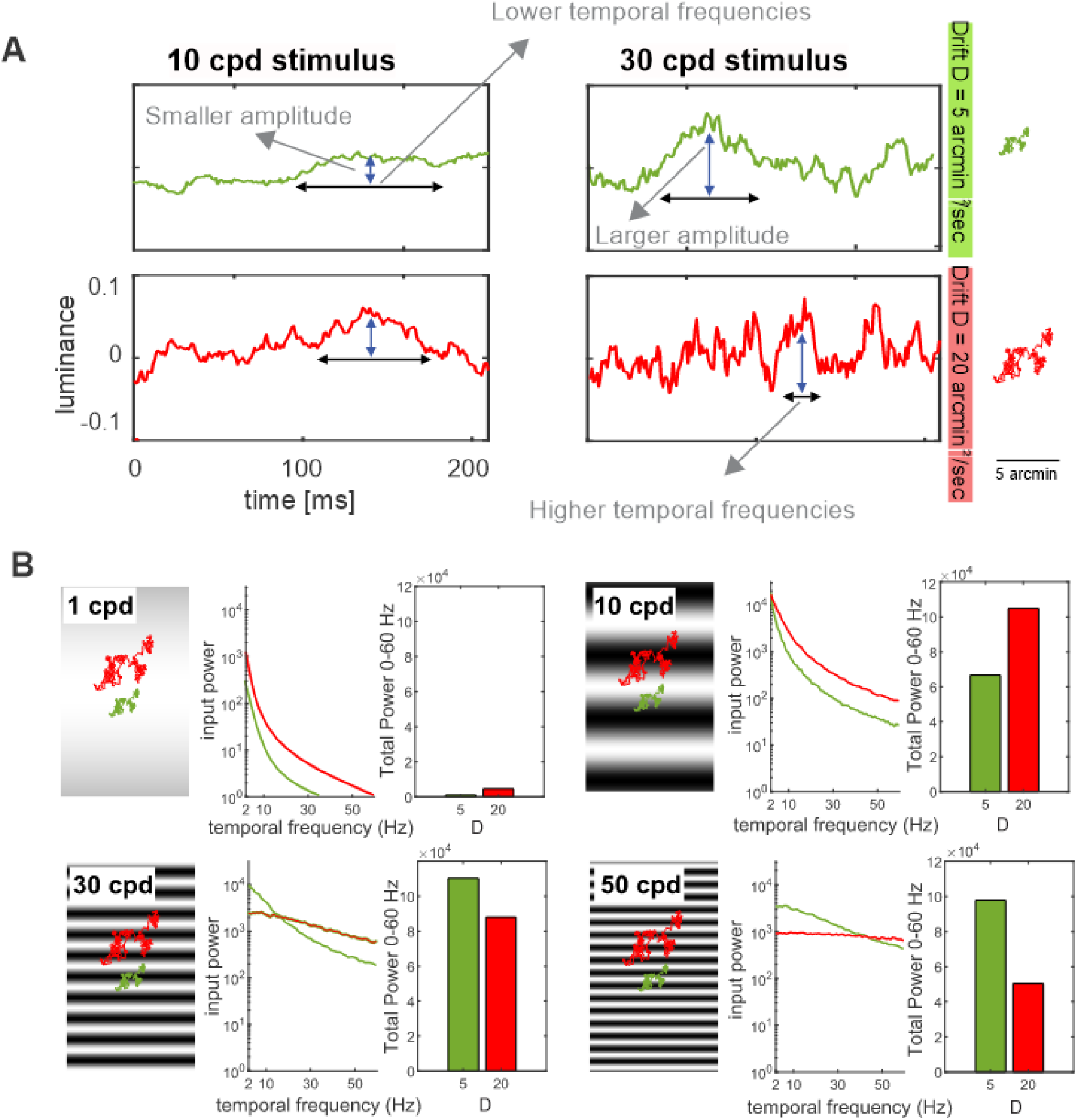
Consequences of ocular drift on visual input signals. (**A**) Luminance modulations resulting from viewing gratings at two spatial frequencies (10 and 30 cycles/deg; separate columns) with the eye drifting by two different amounts (diffusion constant *D* = 5 and 20 arcmin^2^/s; separate rows). Increasing spatial frequency and/or the extent of drift results in larger and faster modulations. (**B**) Power spectra of the luminance modulations delivered by the two types of drifts in *A* when examining stimuli at four spatial frequencies. To provide a measure of the efficacy of these signals in eliciting visual responses, the bar charts show the integral of all power at non-zero temporal frequencies up to 60 Hz, the approximate range of retinal sensitivity. Analyses are based on 1000 simulated drift segments. Note that with a low spatial frequency stimulus, a more diffuse drift generates stronger modulations than a concentrated drift. The opposite occurs with a high spatial frequency stimulus.

These effects are described more quantitatively by the spectral redistributions in Fig. 2*B*. If the spatial frequency of the grating is low (*e*.*g*., 1 cpd), luminance modulations have small amplitudes and are concentrated at low temporal frequencies. In this case, enlarging the drift amplifies modulations without making them too fast, so that a drift with *D* = 20 arcmin/s^2^ yields more power than one with *D* = 5 in the temporal frequency range relevant for retinal neurons. Thus, we would expect a more diffuse drift to be beneficial at low spatial frequencies. As we increase the spatial frequency of the stimulus, however, the temporal bandwidth of the input signals also increases, and progressively more power is shifted toward high temporal frequencies. This effect is accentuated by a broader drift, which depletes power in the useful temporal range of sensitivity, resulting in a less effective driving signal during exposure to high spatial frequency stimuli. For example, for a stimulus at 30 cpd, the sum of all dynamic power up to 60 Hz decreases by 20% when the drift diffusion constant increases from 5 to 20 arcmin/s^2^. Thus, we would expect a more concentrated fixation to be beneficial at high spatial frequencies.

Figure 3 illustrates the hypothesized consequences of ocular drift in two observers, who differ for the degree of acuity afforded by their optics and anatomy (separate panels in the figure). The two curves in each panel represent the strengths of the luminance modulations resulting from drifts with distinct *D*s, each quantified by the total power that the eye motion makes available at non-zero temporal frequencies up to 60 Hz. Because of the effects explained in Fig. 2*B*, the strength of the input signal follows a band-pass function peaking at a critical spatial frequency. Importantly, this critical frequency varies with the amount of drift: the larger the drift—*i*.*e*., the higher its *D*—, the lower the spatial frequency that yields the strongest signal (Fig. 1*C*).

**Figure 3:**
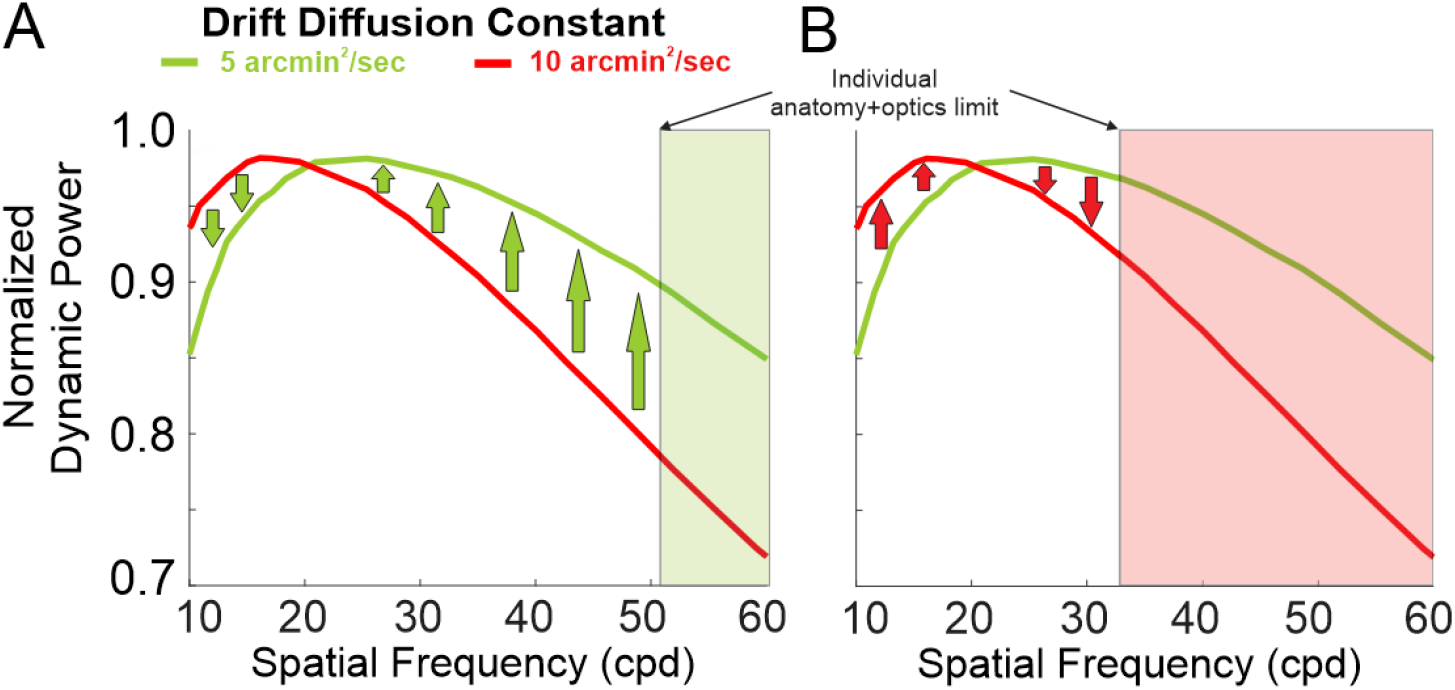
Ocular drift and visual acuity. Predicted consequences of drift modulations in relation to the individual acuity limits afforded by optical, anatomical, and functional characteristics. (**A**) An observer capable of resolving very fine patterns benefits from the increased power at high spatial frequencies delivered by a more concentrated drift. (**B**) In contrast, a larger drift is preferable for an observer with lower acuity, as this motion delivers more power at lower spatial frequencies.

These considerations lead to specific predictions. If observers indeed make use of oculomotor luminance modulations, as supported by a growing body of evidence ^26–28,37^, we would expect a small drift to be beneficial in a subject whose anatomy and optics afford resolution of high spatial frequencies (Fig. 3*A*). This motion increases the critical frequency, shifting power from low to high spatial frequencies in a range that can be detected by this observer’s retinal circuitry. In contrast, we would expect an observer with more limited optics and/or lower density of photoreceptors to be impaired by this redistribution, as much of the resulting power would be beyond his/her acuity limits. In this subject, a larger drift could be more effective, as it yields stronger modulations in a lower spatial frequency range. Therefore, we expect the extent of drift across observers to be positively correlated with acuity, so that the smallest drifts occur in the observers with the highest acuity. In addition, for each individual observer, irrespective of their individual acuity limit, we would expect higher acuity in the trials in which drift remains more concentrated.

To determine whether individual differences in acuity are accompanied by corresponding changes in fixational drifts, we measured visual acuity thresholds in healthy human observers while recording their eye movements. Fixational eye movements were recorded using a high-precision Dual Purkinje Image eye-tracker ^38^ coupled with a system for gaze contingent display control ^39^, an apparatus that enables more accurate localization of the line of sight than standard eyetracking techniques ^40^. Subjects were asked to determine the identity of a briefly presented digit among four possible choices (Fig. 4*A*).

**Figure 4:**
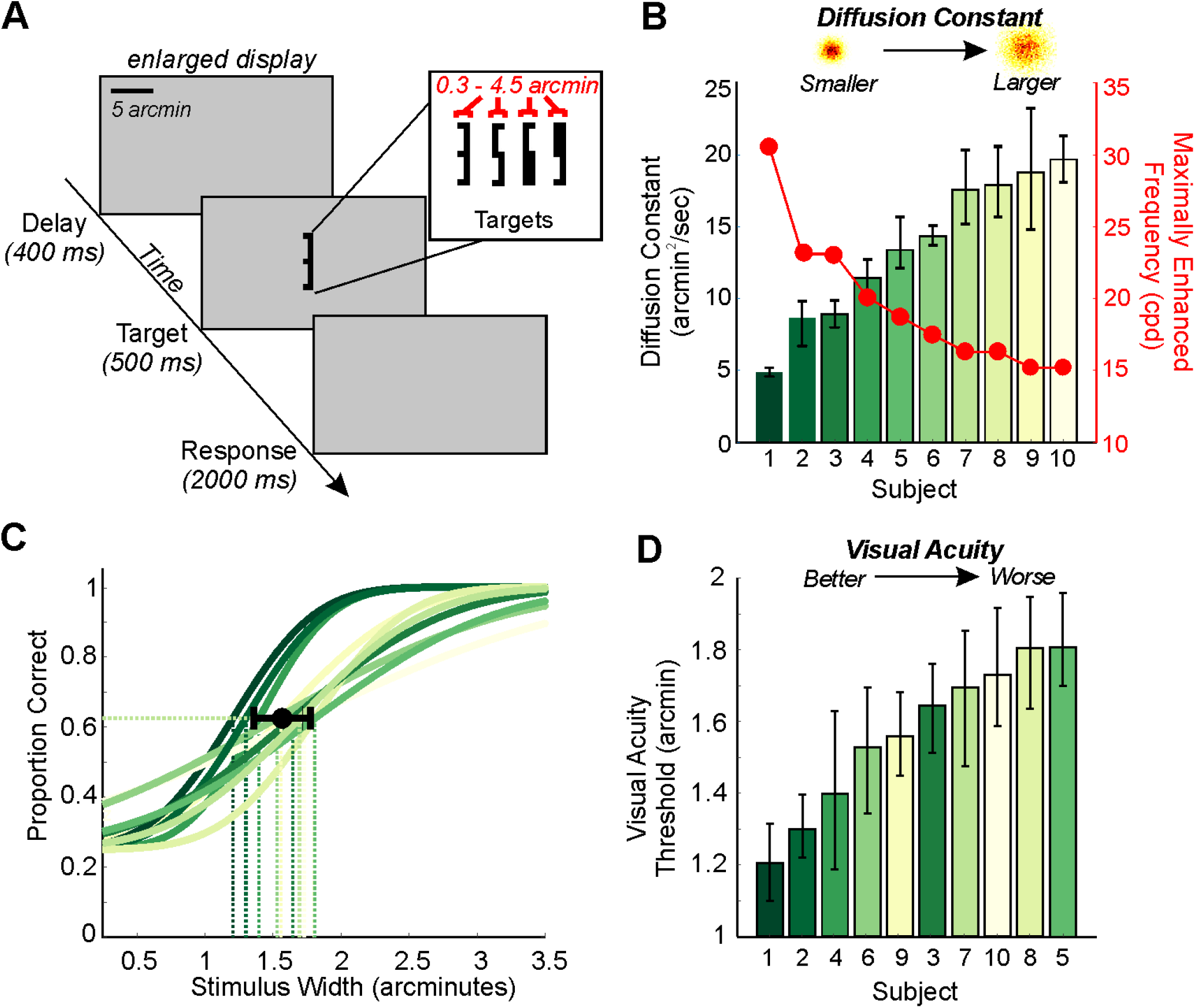
Experimental procedure and behavioral results. (**A**) Subjects were instructed to identify high-acuity stimuli presented at the center of the display for 500 ms. After a brief period of fixation, the target appeared following a delay of 400ms. Stimuli, digits in Pelli font ^41^, varied in size from 0.5 to 4 arcminutes. (**B**) Individual drift diffusion constants (color coded for individual subjects, with 95% bootstrapped confidence intervals), and corresponding critical frequencies (red dots). The illustrations on top represent theoretical 2D distributions of gaze position (red colors indicate higher densities) for smaller and a larger diffusion constants. (**C**) Psychometric fits and acuity thresholds (dashed lines) for single subjects (color coded as in **B** in this and following figures). Curves were fitted on ∼200 trials per subject using a cumulative Gaussian function. The average acuity threshold with ± 1 STD across subjects is also shown (black dot). (**D**) Single subject acuity. Error bars represent bootstrapped 95% confidence intervals.

We first examined the extent of ocular drift variations across healthy emmetropic subjects. We found that ocular drift’s diffusion constant varied greatly across individuals, changing by a factor of ≈4 (from 5 to 20 arcminutes^2^/sec; average ± std, 13.6^*′*^ ± 5.0^*′*^) (Fig. 4*B*). Similar variations in drift diffusion constant were also reported when subjects simply maintained sustained fixation on a marker (ranging from 7 to 27 arcminutes^2^/sec; average ± std, 15.0^*′*^ ± 6.5^*′*^). Therefore, these findings indicate that normal ocular drift changes by a remarkably large amount from one observer to another.

These changes in drift have a profound impact on the spatiotemporal frequency content of the input signal. As shown in figure 3*A*, drifts with a smaller *D* enhance spatial frequencies in a higher range compared to drifts with a larger *D*. Therefore, the variations we observed in diffusion constant across subjects lead to meaningful changes in the frequency content of the retinal input (Fig. 4*B*), emphasizing different frequency ranges. A change in diffusion constant from 5 to 20 arcminutes/sec^2^ led to both a shift of approximately 15 cpd in the maximally enhanced spatial frequency (critical frequency) (Fig. 4*B*), and to a reduction of 64% in power in the 30-60 cpd frequency range. The critical frequency varied across subjects from 15 to 30 cpd (average ± std, 19.6 cpd ± 4.8 cpd). Based on these observations, we predicted that variations in the pattern of ocular drift across observers are associated with variations in visual acuity thresholds.

Acuity thresholds also varied greatly across subjects (Fig. 4*C* and Supplementary Fig. 1). The threshold stimulus width varied between 1.2^*′*^ and 1.8^*′*^ (1.57^*′*^ ± 0.21^*′*^) across observers; a 50% change in size between the smaller and larger measured threshold. Even though all subjects were emmetropic, acuity estimates measured during the initial screening and in the experimental task ranged between 20/20 and 20/12 on the Snellen scale, and screening and task measures were overall consistent (Supplementary Fig. 2).

In line with our prediction, differences in ocular drift across subjects were strongly correlated with corresponding differences in high-acuity thresholds (*p* = 0.017, *r* = 0.728; Fig. 5*A*). More specifically, as one would expect from the characteristics of the luminance signal resulting from drift, the higher the acuity, the smaller the ocular drift diffusion constant. This relationship was observed both when acuity was measured psychophysically and when it was assessed with the Snellen Chart (Supplementary Fig. 3). This significant relationship also holds true when using more intuitive measures of the spatial extent of drift motion than the diffusion constant, such as the fixation span (Supplementary Fig. 4).

**Figure 5:**
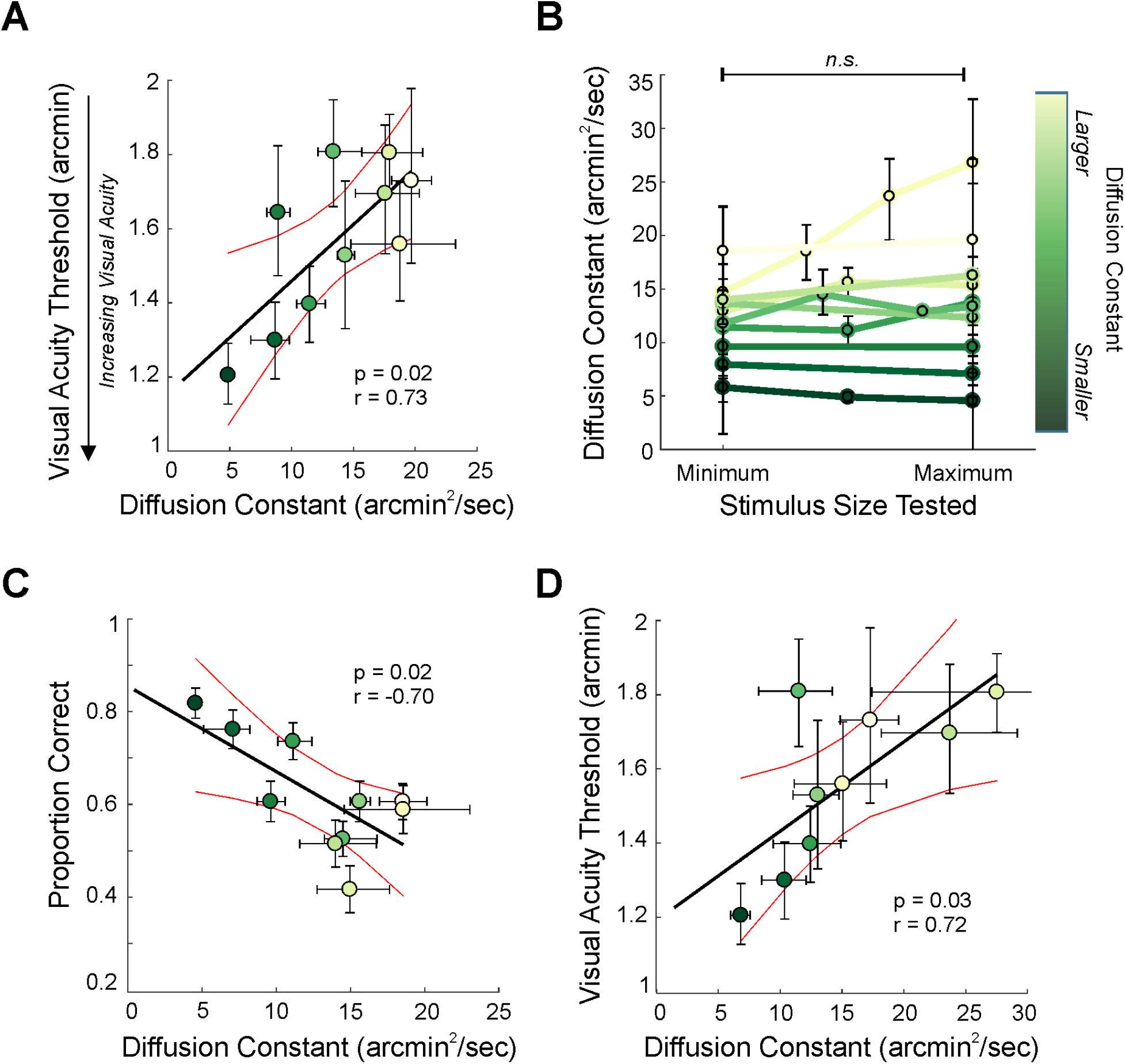
Acuity thresholds and drift characteristics. (**A**) Visual acuity thresholds as a function of diffusion constant in the task. Each point represents an individual subject, error bars are 95% confidence intervals. (**B**) Individual subject diffusion constants as a function of stimulus size. The range of stimulus sizes tested was normalized within each subject. (**C**) Proportion of correct responses for a fixed stimulus size (1.5^*′*^) as a function of drift diffusion constant across subjects. Error bars for proportion correct represent sem. (**D**) Visual acuity thresholds as a function of the diffusion constants measured during sustained fixation on a marker for 9 subjects (note, one subject was removed due to too few fixation trials).

We wondered whether the observed differences in drift characteristics were caused by the stimulus itself. In the experiment, the stimulus changed size following an adaptive procedure to measure acuity thresholds. Since every observer was tested extensively, the stimulus size was near threshold in the majority of trials. This implies that, on average, subjects with worse acuity (higher thresholds) viewed larger stimuli than subjects with better acuity. Thus, a change in ocular drift with stimulus size could, in principle, also explain the observed trend. However, our data do not support this hypothesis. First, drift diffusion constant remained approximately uniform across stimulus sizes (Fig.5*B*, paired t-test *p* = 0.704, comparing diffusion constant for the smallest and the largest stimulus sizes viewed across subjects). With the exception of one subject, the difference in drift diffusion constant between the smallest and the largest stimuli tested was minimal, on average changing 1.8 arcminutes^2^/sec ± 3.9 arcminutes^2^/sec. Second, performance decreased with increasing diffusion constants even when the stimulus size was held fixed (Fig. 5*C, p* = 0.024, *r* = -0.70). Third, the same correlation between drift diffusion constant and acuity was observed when drift was measured independently from the acuity task, *i*.*e*. when subjects simply attempted to maintain steady fixation on a marker (*p*=0.029, *r*=0.72). Remarkably, this shows that acuity can be predicted by measuring drift diffusion constant during a simple sustained fixation task.

Importantly, as shown in Fig.6, despite individual differences in drift diffusion constant, all our subjects were well centered on the target stimuli on average. Stimuli remained within the the central 16^*′*^ x 16^*′*^ region (indicated by orange boxes in Fig. 6) around the center of gaze where acuity is likely uniform ^42,28^, 75.7 ± 12.4% of the viewing time. For each observer, the target was maintained in this region for a duration comparable to or above the amount of time necessary for visual acuity performance to plateau for high contrast stimuli ^43,44^ (mean ± std; 378.5 ms ± 62 ms). Additionally, individual’s average gaze offset from the center of the target was not correlated with acuity (*p* = 0.751, *r* = -0.12).

**Figure 6:**
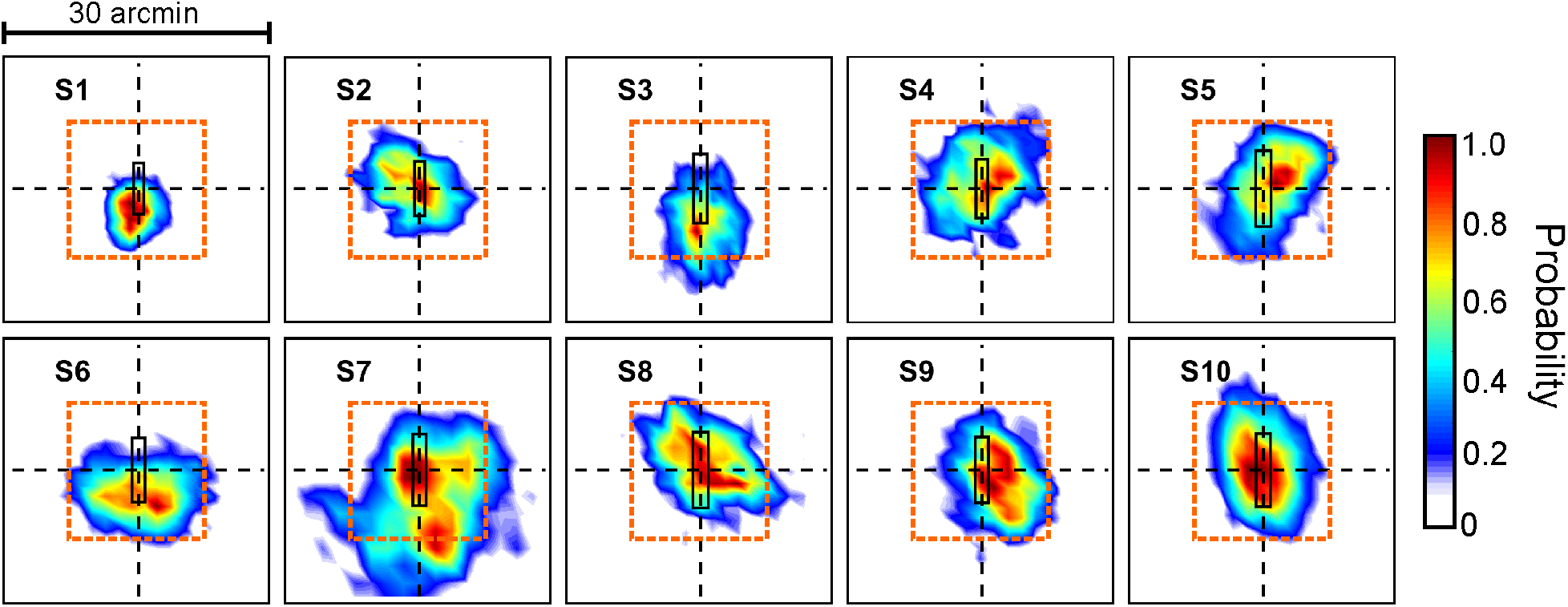
Gaze position during the task. Normalized 2D probability distributions of gaze position during stimulus presentation. Orange boxes delimit the central 16^*′*^x16^*′*^ region. In each graph the central rectangle represents the stimulus size at their threshold acuity.

While the diffusion coefficient provides a good description of the overall amount of drift, it does not capture its directionality. In particular, because of the nature of the stimuli used in the task, which were thin and slender, vertically oriented drifts would introduce stronger luminance modulations compared to horizontally oriented drifts. These influences may also be further amplified by retinal processing ^45–47^. On average, across all trajectories, idiosyncratic biases were visible in the overall drift distributions, as previously reported ^5^ (see supplementary Fig. 5). These biases were, however, small, as drift always moved gaze in all directions by frequently changing course. In keeping with this observation, there was no clear relationship between individual direction bias and acuity, *i*.*e*. subjects with more vertically oriented drifts did not have higher acuity (linear regression, p = 0.21). Additionally, performance in the acuity task did not change when we compared performance between trials with predominantly vertical and horizontal drifts (paired t-test, p = 0.92, Supplementary Fig. 6). This equivalence may be caused by both the small amplitude of biases—possibly not sufficient to influence performance—or the presence in all observers of strong luminance modulations caused by the frequent directional changes of ocular drift. Nevertheless, these results confirm that individual differences in drift direction did not influence our main findings.

To further quantify the impact of ocular drift on acuity, we examined the changes in input luminance introduced by drift. We calculated the total power of these luminance changes for each subject and each digit (given a 1.5^*′*^ width) used in the task (see methods for details). Figure 7*A* shows individual subjects’ performance as a function of the total power conveyed to the retina averaged across the four digits. The overall input power was higher for subjects characterized by higher performance in the task and smaller drift diffusion constants (*p* = 0.017, *r* = 0.728). Performance, with a same size stimulus, decreased by approximately 29% for the subject with the least amount of power. To illustrate how the incoming visual flow to the retina differs for subjects with small vs. large drift diffusion constants, Figure 4*B* shows the power of the stimulus for a theoretical small and a large drift diffusion constant, respectively. A smaller diffusion constant sharpens the edges of the stimuli to a greater extent, making it easier to discriminate across stimuli (see also Supplementary Fig.7 and Supplementary Movie 1).

**Figure 7:**
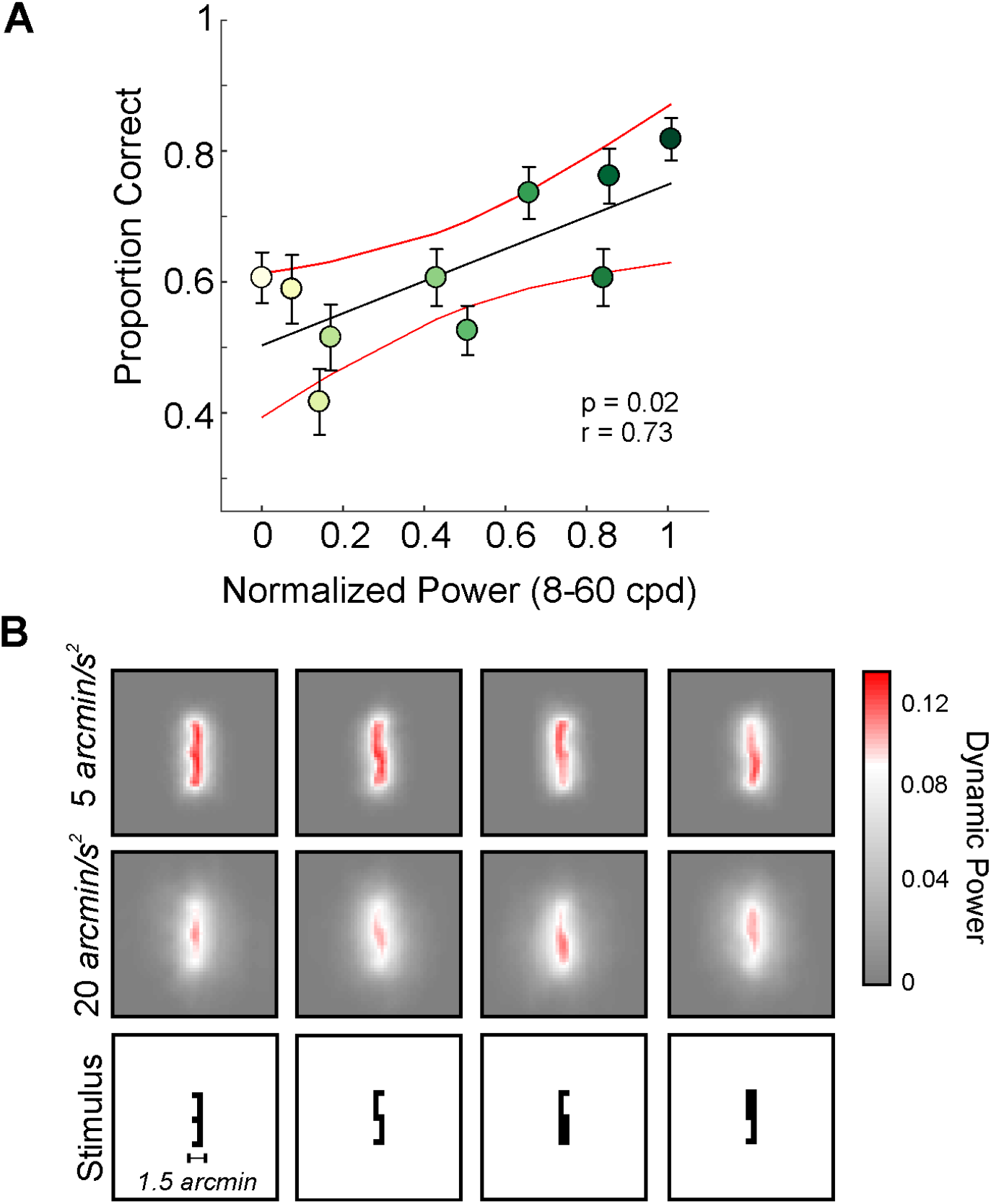
Drift induced modulations of the retinal input during the task. (**A**) Performance in the task as a function of total power conveyed to the retina (averaged across digits 1.5^*′*^ wide). Each circle represents a subject. Error bars represent 95% confidence intervals. (**B**) Power of the retinal input for a theoretical drift with small and large drift diffusion constant respectively (top rows), and the stimulus input (bottom row).

Therefore, differences in the pattern of ocular drift well explain the observed changes in high-acuity vision across subjects; smaller drifts increase the power in the frequency range that is crucial to resolve high-acuity stimuli.

Consideration of the visual input signals resulting from ocular drift (Fig. 3) indicates that an influence of drift amount on performance should also be visible in the data from individual observers. To examine this, we estimated the drift diffusion constant in each single trial, while observers reported on stimuli at threshold. Based on the distribution of *Ds* across trials, we then compared performance between the pools of trials in which the eye moved substantially more or less than the individual subject’s average (trials above the 70^*th*^ percentile vs trials below the 25^*th*^ percentile). If, indeed, ocular drift modulates acuity, we expect trials with a smaller amount of drift to be associated with higher performance in the task. Figure 8 and Supplementary Fig. 8 illustrate that, consistent with these predictions, performance dropped in trials with larger eye motion (paired t-test, *p* = 0.033). This result further demonstrates that acuity is modulated by the specific pattern of ocular drift and that—up to a limit—reducing the amount of drift improves acuity.

**Figure 8:**
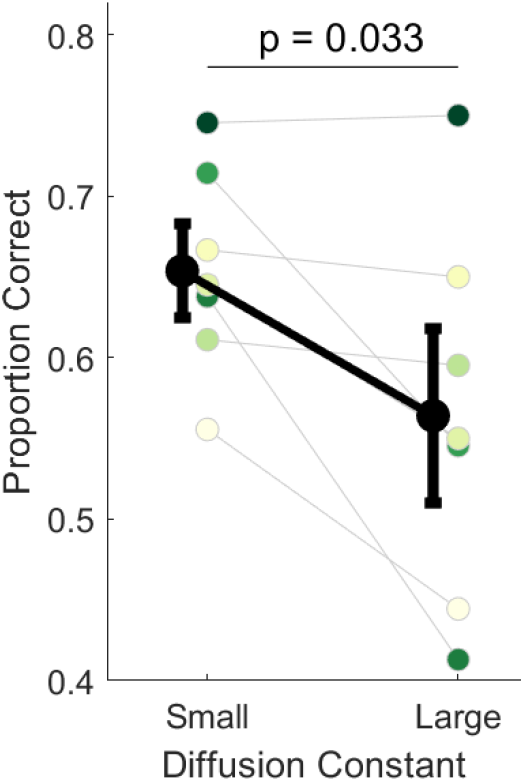
Comparison of performance between trials with different amount of motion. Average and single subject performance in trials in which the eye moved less or more than normal during viewing of threshold stimuli (N=7). Error bars are 95% CI. Colored symbols represent individual subject data, with colors coding their amounts of drift as in Fig. 4*B*. Only subjects with more than 25 trials in each condition were included in the analysis.

## Discussion

The human eyes drift incessantly during fixation, a motion that transforms spatial patterns into temporal modulations on the retina. This transformation maximally redistributes power in a range of spatial frequencies that depends on the amount of drift: the smaller the jitter, the higher the spatial frequency enhanced. Our results show that the idiosyncratic variations in the patterns of ocular drift measured across observers alter considerably the spatial frequency range with strongest oculomotor-induced luminance modulations. We have shown that, as predicted from these considerations, acuity differences across healthy emmetropes are related with variations in individual drift characteristics, so that subjects with highest acuity are also those that drift in a way that emphasizes higher spatial frequencies. Furthermore, our data show that a similar relation can be observed in individual subjects, as the physiological variability of drift influences acuity across trials.

These results are robust; the reported relationship between the amount of ocular drift and acuity is independent from the method used to measure visual acuity. Indeed, we found the same relationship whether using the traditional Snellen chart or a more controlled psychophysical method with stimuli viewed in isolation. Additionally, the same relationship between ocular drift and acuity was also found when ocular drift was measured in a simple oculomotor task that involved no measurement of acuity, subjects simply maintained fixation on a marker. This finding indicates that these results can be reproduced under different conditions. It also shows that in healthy emmetropes—observers with a minimum of 20/20 Snellen acuity without reported ocular pathologies—an informed guess about their individual visual acuity can be obtained from their oculomotor behavior at fixation, and vice versa, their acuity informs about the extent of drift. Our findings go beyond previous work in that they do not simply show that drift is beneficial for fine spatial vision, but that the amount of drift is related to acuity in a specific way. Given that drift can be modulated by the task ^28,48^, it is conceivable that individuals tune their drifts to match the optical/anatomical characteristics so to optimize the extraction of fine spatial information on the basis of the capabilities afforded by their eyes.

Given that the luminance modulations from progressively smaller drifts emphasize increasingly higher spatial frequencies, one may wonder whether smaller drifts than those measured in our experiments could further improve acuity. One possibility is that achieving finer drifts may not be feasible because of limitations in oculomotor control. Yet, even if the oculomotor system is capable of reducing drift further, this may not be beneficial. Smaller drifts will enhance spatial frequencies beyond 60 cycles/deg, a range that the retina would not be able to resolve. This enhancement occurs simultaneously with an attenuation in the strength of the signal at lower spatial frequencies (see Fig.3*A*), effectively yielding a less contrasted input in the visible frequency range. Therefore, further reducing drift motion may not be advantageous, particularly considering that visual stimuli normally possess both low and high spatial frequencies, the relative importance of which depends on the task.

Related to these considerations, the finding that smaller drift motion is associated with higher acuity may lead to conclude that no motion at all would be the most beneficial condition for high-acuity vision. This however is not the case. Multiple studies have already shown how retinal stabilization decreases performance in high-acuity tasks ^26,28,6,27^. These findings are consistent with the response characteristics of neurons in the early stages of the visual system. In the absence of other forms of temporal modulations, eliminating drift motion entirely will concentrate all power of the retinal input at 0 Hz. This effect is expected to lead to an overall decreased neural response, as neurons are mostly sensitive to non-zero temporal frequencies ^32,49^.

The results described here prompt a number of questions. Does drift vary over the course of the lifespan and with changes in the refractive error? Little is known on this. However, our results showing a clear relationship between drift and acuity and the ability of drift to modulate acuity, suggest that drift may change over an individual’s lifetime. Interestingly, acuity and fixational stability have been shown to be worse during early childhood ^50–52^, further suggesting that drift may change based on the varying optical/anatomical constraints from childhood to adulthood. Another open question pertains to the relationship between ocular drift and foveal anatomy. During the course of 500 ms of fixation on average drift moves a point stimulus over ≈35 photoreceptors, assuming an average photoreceptor size of 0.5’ at the foveal center. A change in *D* by a factor of 4 leads to fourfold increment in the number of cones being stimulated. Crucially, cone density varies greatly across subjects ^53^, therefore the same drift pattern may have different impacts depending on an individual’s cone density.

Our findings of how acuity is related to the physiological variability in drift suggest that subjects can potentially improve their visual acuity by changing their drift pattern. Previous work has shown task-dependent changes in drift characteristics ^54^. Recent work has also reported that individuals express differences in ocular drift movements across various conditions, especially when looking at different types of stimuli ^28,48^, and when the head is free ^55^. While there is evidence indicating that highly trained subjects are characterized by more stable fixation ^5^, and drift can be actively modulated depending on the viewing condition ^28^, it remains unclear if ocular drift can change with training and experience. In light of the findings reported here, understanding if ocular drift can be shaped through training has important clinical and practical implications, potentially allowing for improvements in fine spatial vision.

In sum, we have shown that ocular drift is an important component in the equation explaining humans’ ability to achieve visual acuity. These findings raise important questions on the visuomotor strategies present during vision of spatial detail and the principles controlling fixational drift. A stimulating hypothesis is that healthy emmetropes optimize eye motion based on the constraints imposed by optics and anatomy to yield the strongest signal in the spatial frequency range afforded by their individual eye characteristics. Further research is needed to investigate this hypothesis and elucidate the interplay between drift, optics and retinal anatomy.

## Methods and Procedures

### Observers

10 adult observers with normal vision participated in this study. 9 naive subjects, and 1 experienced observer (Subject 2) who is an author. Subjects ranged in gender (3 males, 7 females) and age from 18 to 25 years old. During screening, subjects were asked to perform a Snellen Acuity chart test. All subjects except for the author were emmetropic and required no correction to reach a minimum of 20/20 Snellen acuity. The author wore corrective contact lenses. At the end of the study, the Reichert RK600 Autorefractor was used to estimate subject’s refraction. 3/9 emmetropic subjects were able to perform this measure and showed no significant spherical (average -0.6D) or astigmatic (less -0.25 cylinder) correction was needed. Subjects for whom we did not evaluate their refraction did not have a history of reported astigmatism. This research study was approved by the University of Rochester’s Research Subjects Review Board. Subjects were invited for an initial screening session, which involved a thorough explanation of the experiment, as well as a detailed review of the materials in the consent form. After the subject understood the information in the consent form and verbally agreed to participate in the study, informed consent was obtained and documented.

### Stimuli and Apparatus

Stimuli were presented monocularly to the right eye while the left eye was patched. Eye movements were recorded with high precision either by means of a Generation 6 Dual Purkinje Image (DPI) eye tracker (Fourward Technologies), with a 1kHz sampling rate ^56,38^, or by means of a custom-made digital Dual Purkinje Image (dDPI) eye tracker, with a sampling rate of 340Hz ^57^. Both systems have an internal noise well below 1^*′*^, and a spatial resolution of at least 1^*′*^ ^56,38^. To reduce noise and achieve higher precision in the eyetracking signal, the head was immobilized by means of a dental-imprint bite bar and head-holder. Stimuli were shown on a LCD monitor (ASUS PG258Q), with a vertical refresh rate of 200Hz, and a spatial resolution of 1920 × 1080 pixels. The monitor was between 3 or 5 meters away from the observer (1 pixel = 0.25^*′*^ and 1 pixel = 0.19 ^*′*^, respectively).

Stimuli consisted of 3,5,6 and 9 digits from the Pelli number-font ^41^. These targets were presented in isolation at the center of the display (emphasized by presenting 4 arches in the periphery centered on the stimulus) for 500 ms. Conveniently, these stimuli also allow us to compare performance in isolation with performance under crowded conditions in the fovea, overcoming issues that would normally arise when using traditional optotypes ^41^. Stimuli consisted of black font presented on a gray background. Stimuli were rendered by means of EyeRIS ^39^, a custom-developed system allowing flexible gaze-contingent display control. This system acquires eye movement signals from the eye-tracker, processes them in real time and, if necessary, updates the stimulus on the display according to the desired combination of estimated oculomotor variables.

### Experimental Paradigm

Data were collected by means of multiple experimental sessions. Each session lasted approximately 1 hour, and each subject completed on average 5 sessions. Every session started with preliminary setup operations that lasted a few minutes, involving comfortably positioning the observer in the apparatus, tuning the eye tracker for optimal performance, and executing a two-step gaze-contingent calibration procedure to map the eye tracker’s output into visual angle. This procedure improves localization of the preferred retinal locus of fixation by approximately one order of magnitude over standard methods ^40^. In the first phase (automatic calibration), observers sequentially fixated on each of the nine points of a 3×3 grid, as it is standard in oculomotor experiments. Points in the grid were 1 or 1.25^*°*^ apart from each another on the horizontal axes, and 40 or 50^*′*^ on the vertical axes (varying based on screen distance). In the second phase (manual calibration), observers confirmed or refined the mapping given by the automatic calibration by fixating again on each of the nine points of the grid while the location of the line of sight estimated on the basis of the automatic calibration was displayed in real time on the screen. Observers used a joypad to fine-tune the predicted gaze location if necessary. The manual calibration procedure was repeated for the central position before each trial to compensate for possible microscopic head movements and system drift that may occur even on a bite bar.

Once the subjects initiated the trial, a brief 10×10^*′*^ fixation point was presented at the center of the screen to clearly identify the location where the stimuli would appear. A 400ms delay period followed the blank screen to avoid any aftereffects from the fixation point. The target was then presented. Subjects were asked to identify the stimulus, choosing among 4 possible digits, by pressing a button on a remote controller. Trials in which subjects were required to maintain fixation on a 10×10^*′*^ marker (fixation trials) at the center of the display were presented once every 30-50 task trials. In fixation trials subjects were instructed to maintain fixation for 2-5 seconds.

Target acuity was determined by following the Parametric Estimation by Sequential Testing (PEST) procedure ^58^, according to which, size would change online based on subject performance.

## Data Analysis

### Eye Movements

Eye movements were categorized into two main groups: saccades (including microsaccades) and ocular drift. Ocular motion in between saccades was defined as drift. Classification of these eye movements was first performed automatically, then thoroughly reviewed by an expert experimenter. Trials containing saccades, blinks and/or bad tracking during stimulus presentation were removed. Furthermore, trials in which subjects did not respond or gaze was more than 30^*′*^ from the center fixation point at the beginning of the trial were also removed (on average 1.3% ± 0.58% of the total trials). We examined ocular drift at regime, far from saccades or blinks. Drift acted as a stationary process in these intervals and its speed did not change during the course of fixation (Supplementary Fig. 9).

### Estimation of Acuity Thresholds

Visual acuity threshold, *i*.*e*. the minimum stimulus width required to perform reliably above chance level (62.5% correct,with a 25% chance level), was determined using a cumulative Gaussian psychometric function ^59^. Better acuity corresponds to lower threshold values.

### Drift Diffusion Constant and Power Spectrum analysis

Ocular drift was characterized with the diffusion constant. Drift diffusion constant can be described as the rate at which the variance of the drift changes over different time delays. If we assume that drift behaves like Brownian motion in the short time of a fixation, the probability of gaze position varies with time as:

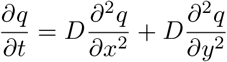

Where *q* is the probability of gaze being at a given location at time t and *D* is the diffusion constant which regulates how quickly gaze moves away from the initial position. Diffusion constant was determined based on the empirical average 2D eye displacement as a function of time for time delays ranging from 50 to 256 ms. Eye displacement data was then fitted with a linear regression and the diffusion constant was calculated as the slope of the fitted line divided by 4. Deviations from the fit were minimal for all subjects (Supplementary Fig. 10). The diffusion constant (*D*) was therefore estimated as following:

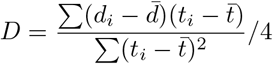

Where *d*= ⟨*x*^2^+*y*^2^⟩ represents the displacement square and t is the temporal interval over which the displacement square is calculated.

As eye drift can be well modeled by Brownian motion ^33^, the power spectrum (*Q*) of drift can then be directly derived from its diffusion constant as:

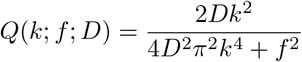

Where *k*^2^ is 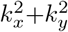 and *k* and *f* are the spatial and the temporal frequencies respectively.

The exact critical frequency (*CF*), which depends on the statistics of ocular drift, can be derived with the following equation:

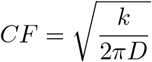

As a result, the larger the drift diffusion constant, the lower the CF.

The drift dynamic power was calculated by summing across all non-zero temporal frequencies of the drift power spectrum (see Fig. 7*A*) multiplied by the power spectrum of the external stimulus (for stimuli of 1.5^*′*^ in width). To quantify individual differences, we summed the resulting power spectrum across all spatial frequencies between 8-60 cpd, *i*.*e*., the most informative range of spatial frequencies for discriminating across stimuli. This operation was performed for each subject and for all digits used in the task. Since the total power yielded varied greatly across digits used in the task, In Figure 7*B*, the plotted power was normalized per each digit across subjects. The normalized power was then averaged per each subject.

All data and MATLAB scripts used to create the figures in the manuscript have been uploaded onto the Open Science Framework repository.

## ACKNOWLEDGMENTS

This work was supported by the National Institutes of Health grants R01EY029788-01 (to M.P.), R01EY18363 (to M.R.) and F31EY029565 (to J.I.). We thank Greg DeAngelis, Jude Mitchell and Florian Jaeger for helpful comments and discussion during the course of this research.

## Notes

### Competing Interest Statement

The authors have declared no competing interest.

### Summary of Updates

Changes made to reflect article submitted and accepted for publication after peer-review.

## References

1. Rovner, B.W. and Casten, R.J. Activity loss and depression in age-related macular degeneration. American Journal of Geriatric Psychiatry, 10(3):305–310, 2002.

2. Hassell, J.B., Lamoureux, E.L., and Keeffe, J.E. Impact of age related macular degeneration on quality of life. British Journal of Ophthalmology, 90(5):593–596, 2006.

3. Kolb, B., Forgie, M., Gibb, R., Gorny, G., and Rowntree, S. Age, experience and the changing brain. Neuroscience & Biobehavioral Reviews, 22(2):143–159, 1998.

4. Steinman, R.M. and Collewijn, H. Binocular retinal image motion during active head rotation. Vision Research, 20(5):415–429, 1980.

5. Cherici, C., Kuang, X., Poletti, M., and Rucci, M. Precision of sustained fixation in trained and untrained observers. Journal of Vision, 12(6):1–16, 2012.

6. Rucci, M. and Victor, J.D. The unsteady eye: an information-processing stage, not a bug. Trends in Neurosciences, 38(4):195–206, 2015.

7. Barlow, H.B. Eye movements during fixation. The Journal of Physiology, 116:290–306, 1952.

8. Burak, Y., Rokni, U., Meister, M., and Sompolinsky, H. Bayesian model of dynamic image stabilization in the visual system. 107:19525–19530, 2010.

9. Anderson, A.G., Ratnam, K., Roorda, A., and Olshausen, B.A. High-acuity vision from retinal image motion. Journal of Vision, 20(7):1–19, 2020.

10. Packer, O. and Williams, D.R. Blurring by fixational eye movements. Vision Research, 32(10):1931–1939, 1992.

11. Rossi, E.A. and Roorda, A. The relationship between visual resolution and cone spacing in the human fovea. Nature Neuroscience, 13(2):156–157, 2010.

12. Green, D.G. Regional variations in the visual acuity for interference fringes on the retina. The Journal of Physiology, 207(2):351–356, 1970.

13. Williams, D.R. and Coletta, N.J. Cone spacing and the visual resolution limit. JOSA A, 4(8):1514–1523, 1987.

14. Thibos, L., Cheney, F., and Walsh, D. Retinal limits to the detection and resolution of gratings. JOSA A, 4 (8):1524–1529, 1987.

15. Marcos, S. and Navarro, R. Determination of the foveal cone spacing by ocular speckle interferometry: limiting factors and acuity predictions. JOSA A, 14(4):731–740, 1997.

16. Hering, E. Ü ber die grenzen der sehschärfe. Ber. math.-phys. Cl. D. königl. Sächs. Gesell. Wiss. Leipzig, pages 16–24, 1899.

17. Arend, L.E. Spatial differential and integral operations in human vision: Implications of stabilized retinal image fading. Psychological Review, 80(5):374–395, 1973.

18. Averill, H.I. and Weymouth, F.W. Visual perception and the retinal mosaic. II. The influence of eye movements on the displacement threshold. Psychological Review, 5:147–176, 1925.

19. Marshall, W.H. and Talbot, S.A. Recent evidence for neural mechanisms in vision leading to a general theory of sensory acuity. 7:117–164, 1942.

20. Steinman, R.M. and Levinson, J.Z. The role of eye movement in the detection of contrast and spatial detail. Eye Movements and Their Role in Visual and Cognitive Processes, 4:115–212, 1990.

21. Ahissar, E. and Arieli, A. Figuring space by time. 32(2):185–201, 2001.

22. Watson, A.B. and Turano, K. The optimal motion stimulus. Vision research, 35(3):325–336, 1995.

23. Tulunay-Keesey, U. and Jones, R.M. The effect of micromovements of the eye and exposure duration on contrast sensitivity. Vision Res., 16(5):481–488, 1976.

24. Riggs, L.A., Ratliff, F., Cornsweet, J.C., and Cornsweet, T.N. The disappearance of steadily fixated visual test objects. Journal of the Optical Society of America, 43(6):495–501, 1953.

25. Barlow, H.B. Possible principles underlying the transformations of sensory messages. In Rosenblith, W.A., editor, Sensory Communication, pages 217–234. MIT Press, Cambridge, MA, 1961.

26. Rucci, M., Iovin, R., Poletti, M., and Santini, F. Miniature eye movements enhance fine spatial detail. Nature, 447(7146):852–855, 2007.

27. Ratnam, K., Domdei, N., Harmening, W.M., and Roorda, A. Benefits of retinal image motion at the limits of spatial vision. Journal of Vision, 17(1):1–11, 2017.

28. Intoy, J. and Rucci, M. Finely tuned eye movements enhance visual acuity. Nature Communications, 11(1): 1–11, 2020.

29. Attneave, F. Some informational aspects of visual perception. Psychological Review, 61(3):183, 1954.

30. Ölveczky, B.P., Baccus, S.A., and Meister, M. Segregation of object and background motion in the retina. Nature, 423(6938):401–408, 2003.

31. Rucci, M., Edelman, G.M., and Wray, J. Modeling LGN responses during free-viewing: A possible role of microscopic eye movements in the refinement of cortical orientation selectivity. 20(12):4708–4720, 2000.

32. Purpura, K., Tranchina, D., Kaplan, E., and Shapley, R.M. Light adaptation in the primate retina: Analysis of changes in gain and dynamics of monkey retinal ganglion cells. Visual Neuroscience, 4:75–93, 1990.

33. Kuang, X., Poletti, M., Victor, J.D., and Rucci, M. Temporal encoding of spatial information during active visual fixation. Current Biology, 22(6):510–514, 2012.

34. Campbell, F. and Gubisch, R. Optical quality of the human eye. The Journal of Physiology, 186(3):558–578, 1966.

35. Engbert, R., Mergenthaler, K., Sinn, P., and Pikovsky, A. An integrated model of fixational eye movements and microsaccades. Proceedings of the National Academy of Sciences of the United States of America, 108: 765–770, 2011.

36. Pitkow, X., Sompolinsky, H., and Meister, M. A neural computation for visual acuity in the presence of eye movements. PLoS Biology, 5(12):e331, 2007.

37. Anderson, T.J. and MacAskill, M.R. Eye movements in patients with neurodegenerative disorders. Nature Reviews Neurology, 9(2):74–85, 2013.

38. Ko, H.k., Snodderly, D.M., and Poletti, M. Eye movements between saccades: Measuring ocular drift and tremor. Vision Research, 122:93–104, 2016.

39. Santini, F., Redner, G., Iovin, R., and Rucci, M. Eyeris: a general-purpose system for eye-movement-contingent display control. Behavior Research Methods, 39(3):350–364, 2007.

40. Poletti, M. and Rucci, M. A compact field guide to the study of microsaccades: Challenges and functions. Vision Research, 118:83–97, 2016.

41. Pelli, D.G., Waugh, S.J., Martelli, M., Crutch, S.J., Primativo, S., Yong, K.X., Rhodes, M., Yee, K., Wu, X., Famira, H.F., and Yiltiz, H. A clinical test for visual crowding. F1000Research, 5(81):1–20, 2016.

42. Poletti, M., Listorti, C., and Rucci, M. Microscopic eye movements compensate for nonhomogeneous vision within the fovea. Current Biology, 23(17):1691–1695, 2013.

43. Baron, W.S. and Westheimer, G. Visual acuity as a function of exposure duration. JOSA, 63(2):212–219, 1973.

44. McAnany, J.J. The effect of exposure duration on visual acuity for letter optotypes and gratings. Vision Research, 105:86–91, 2014.

45. Wool, L.E., Crook, J.D., Troy, J.B., Packer, O.S., Zaidi, Q., and Dacey, D.M. Nonselective wiring accounts for red-green opponency in midget ganglion cells of the primate retina. Journal of Neuroscience, 38(6): 1520–1540, 2018.

46. Sachtler, W. and Zaidi, Q. Visual processing of motion boundaries. Vision Research, 35(6):807–826, 1995.

47. Ennis, R., Cao, D., Lee, B.B., and Zaidi, Q. Eye movements and the neural basis of context effects on visual sensitivity. Journal of Neuroscience, 34(24):8119–8129, 2014.

48. Shelchkova, N., Tang, C., and Poletti, M. Task-driven visual exploration at the foveal scale. Proceedings of the National Academy of Sciences, 116(12):5811–5818, 2019.

49. Benardete, E.A. and Kaplan, E. The dynamics of primate M retinal ganglion cells. Visual Neuroscience, 16 (2):355–368, 1999.

50. Morad, Y., Werker, E., and Nemet, P. Visual acuity tests using chart, line, and single optotype in healthy and amblyopic children. Journal of American Association for Pediatric Ophthalmology and Strabismus, 3(2): 94–97, 1999.

51. Dayton, G.O., Jones, M.H., Aiu, P., Rawson, R.A., Steele, B., and Rose, M. Developmental study of coordinated eye movements in the human infant: I. visual acuity in the newborn human: A study based on induced optokinetic nystagmus recorded by electro-oculography. Archives of Ophthalmology, 71(6):865–870, 1964.

52. Kowler, E. and Martins, A.J. Eye movements of preschool children. Science, 215(4535):997–999, 1982.

53. Curcio, C.A. and Allen, K.A. Topography of ganglion cells in human retina. Journal of Comparative Neurology, 300(1):5–25, 1990.

54. Epelboim, J. and Kowler, E. Slow control with eccentric targets: evidence against a position-corrective model. Vision Research, 33(3):361–380, 1993.

55. Poletti, M., Aytekin, M., and Rucci, M. Head-eye coordination at a microscopic scale. Current Biology, 25 (24):3253–3259, 2015.

56. Crane, H.D. and Steele, C.M. Generation-v dual-purkinje-image eyetracker. Applied Optics, 24(4):527–537, 1985.

57. Rucci, M., Wu, R., and Zhao, Z. System and method for real-time high-resolution eye-tracking. U.S. Patent 11003244, May 2021.

58. Taylor, M.M. and Creelman, C.D. Pest: Efficient estimates on probability functions. The Journal of the Acoustical Society of America, 41(4A):782–787, 1967.

59. Wichmann, F.A. and Hill, N.J. The psychometric function: II. bootstrap-based confidence intervals and sampling. Perception & Psychophysics, 63(8):1314–1329, 2001.

