## SupplementaryMaterial for "Eye drift during fixation predicts visual acuity"

### Individual differences in eye drift predict visual acuity

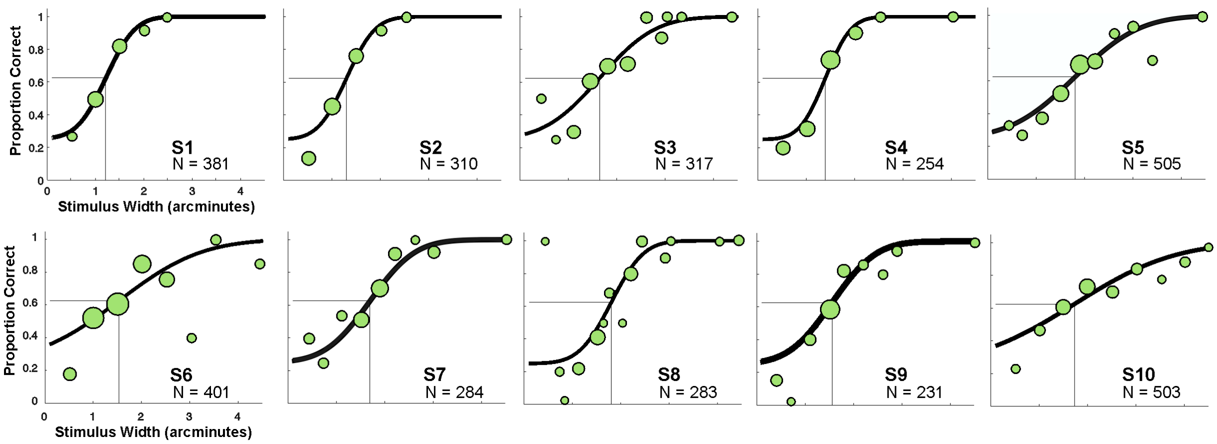


**Fig. S 1**: **Individual psychometric fits for each participant**. The size of points in individual graphs is proportional to the number of trials per stimulus width tested. The total number of trials (N) per subject are also shown in the graphs.

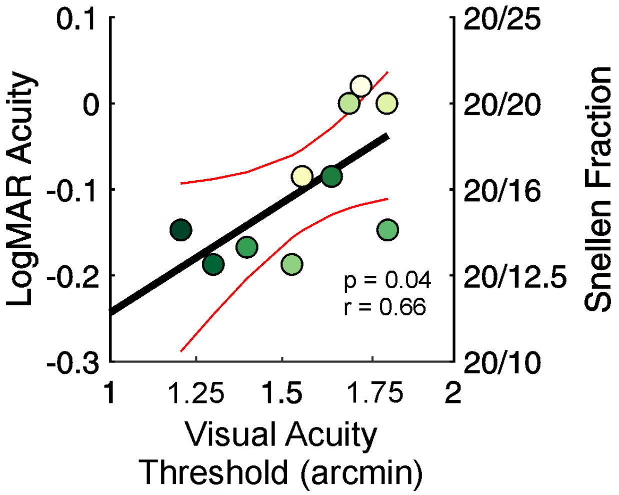


**Fig. S 2**: **Alternative visual-acuity measures**. Traditional visual acuity measured using a Snellen eye chart as a function of acuity measured in our task. Snellen measures are based on one assessment per subject. Each color represents a different subject.

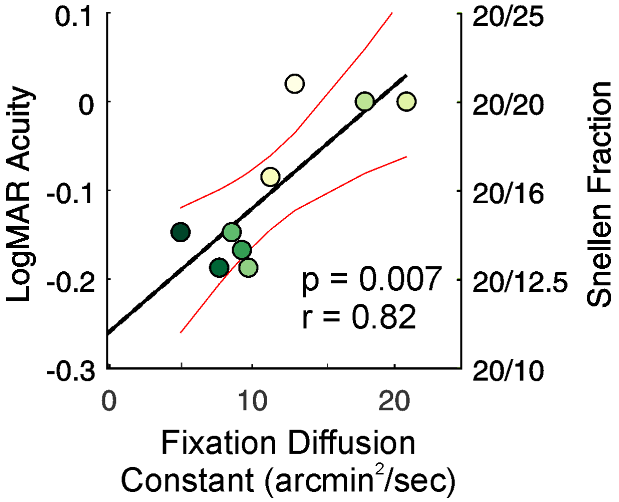


**Fig. S 3**: **The relationship between Snellen acuity and diffusion constant**. Traditional visual acuity measured using a Snellen eye chart as a function of diffusion constant measured during fixation.

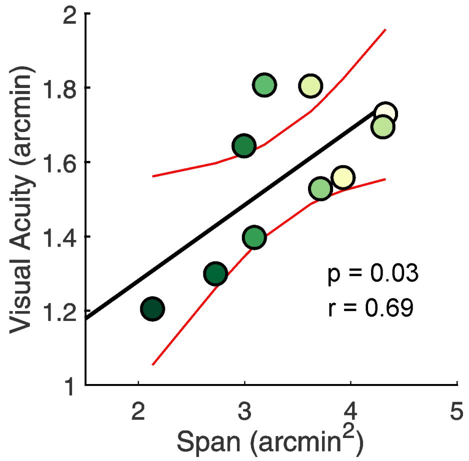


**Fig. S 4**: **Visual acuity measured in the task as a function of ocular drift span**. Drift span is defined as the radius of the smallest circle encompassing drift trajectory. Each color represents a different subject.

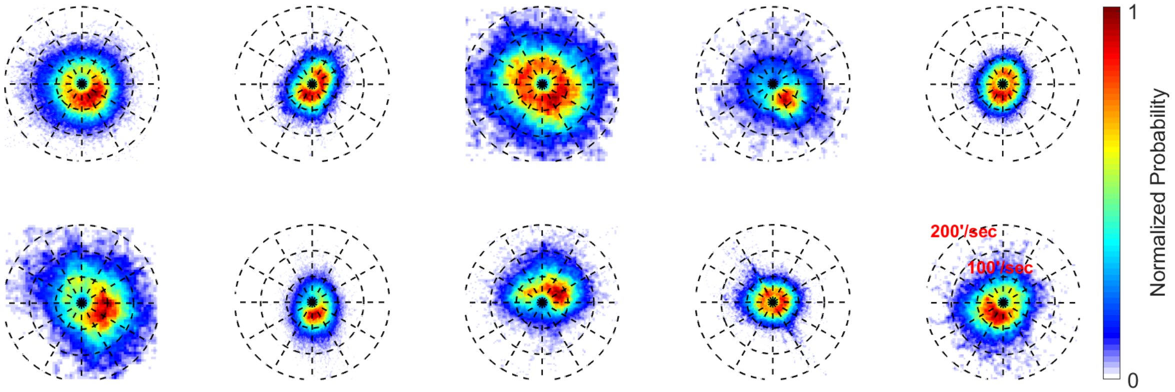


**Fig. S 5**: **Drift Velocity**. 2D normalized probability distributions of instantaneous drift velocity for individual observes. Subjects are ordered from highest (top left) to lowest acuity (bottom right).

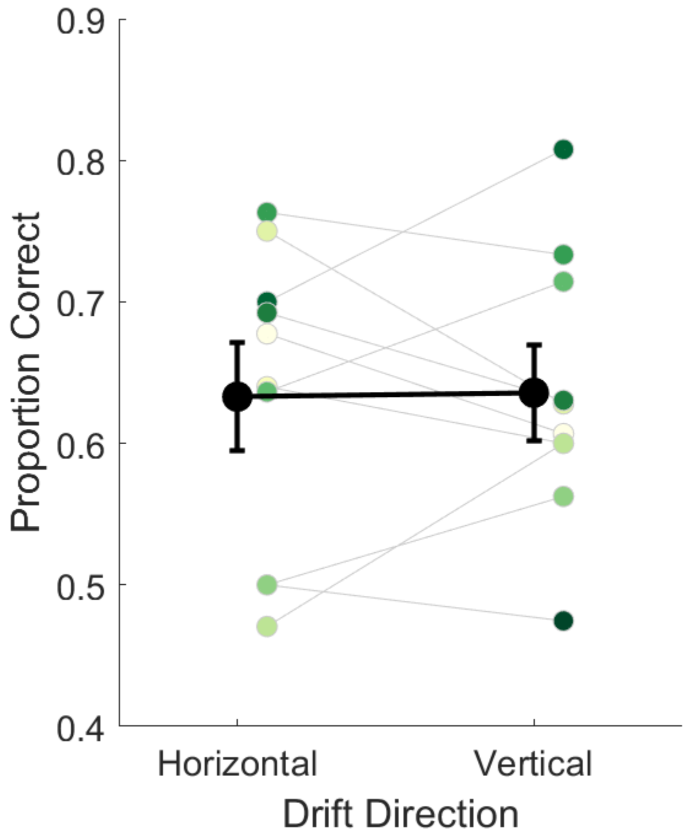


**Fig. S 6**: **Drift direction and performance in the acuity task**. Proportion of correct responses in the acuity task with a fixed stimulus size at threshold for trials with predominantly vertically/horizontally oriented drifts. Black solid lines indicate average performance. Single subjects performances are also shown. Error bars represent 95%CI.

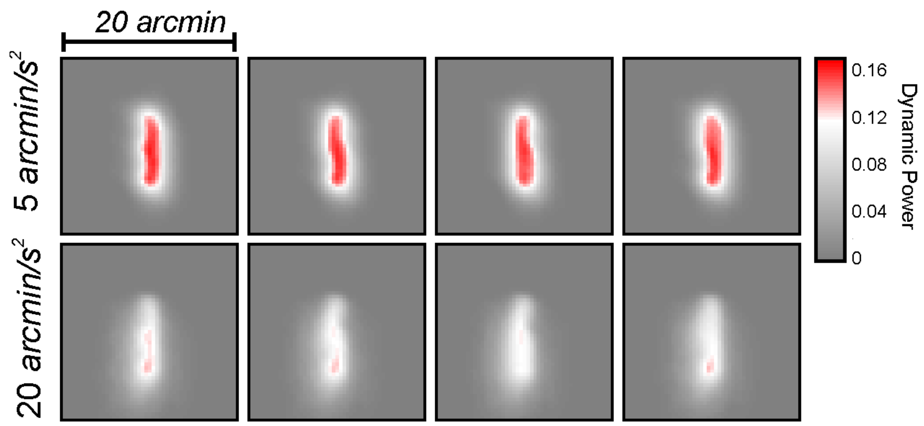


**Fig. S 7**: **Changes in power for differing visual input from eye motion**. Total power of the visual input calculated using eye traces of a subject with a diffusion constant of 5 (top row) and 19 (bottom row) arcmin$^2$/sec, respectively.

**Movie S 1**: **Changes of power in time for different eye motion**. A reconstruction of the drift-induced temporal modulations impinging onto the retina during a fixation period for a drift with a diffusion constant of 5 and 20 $\text{arcmin}^2/\text{sec}$, respectively.

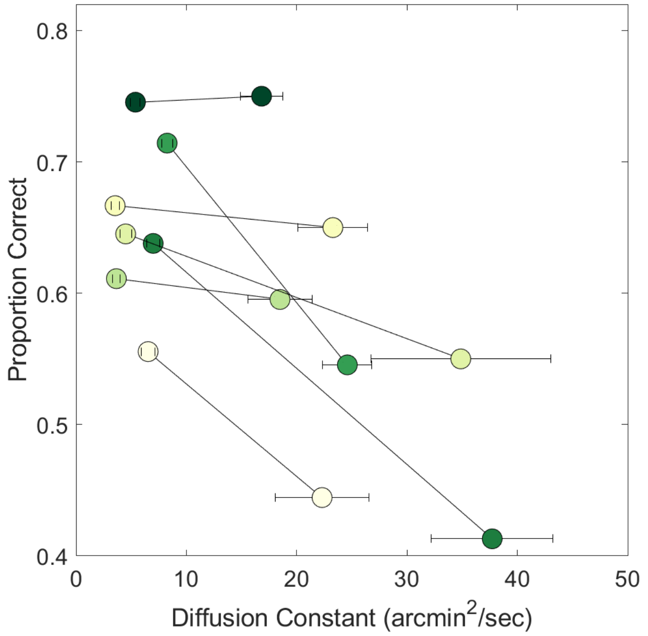


**Fig. S 8**: **Drift diffusion constant and performance in the acuity task.** Individual subject's performance in trials with smaller and larger drift $D$ with respect to the average. Each color represents a different subject. Error bars are 95% CI.

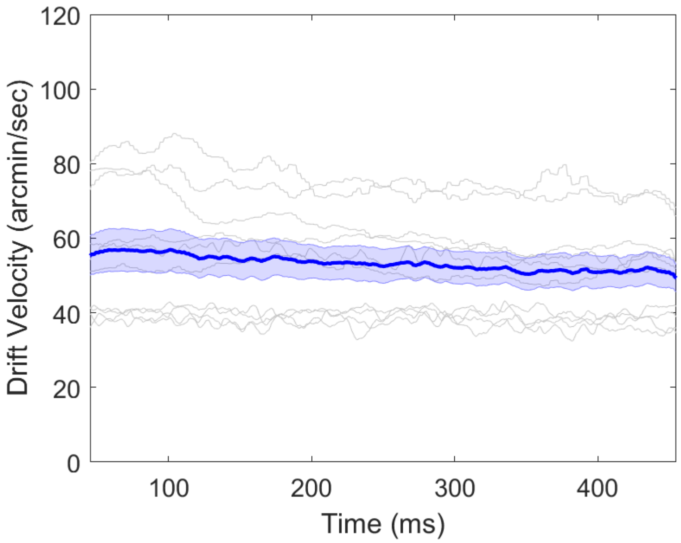


**Fig. S 9**: **Average instantaneous drift speed as a function of time during stimulus presentation.** Shaded regions represent SEM. Single subjects are shown in light gray. Speed was estimated on ocular drift traces filtered with a third order Savitzky-Golay filter with 41 frame length.

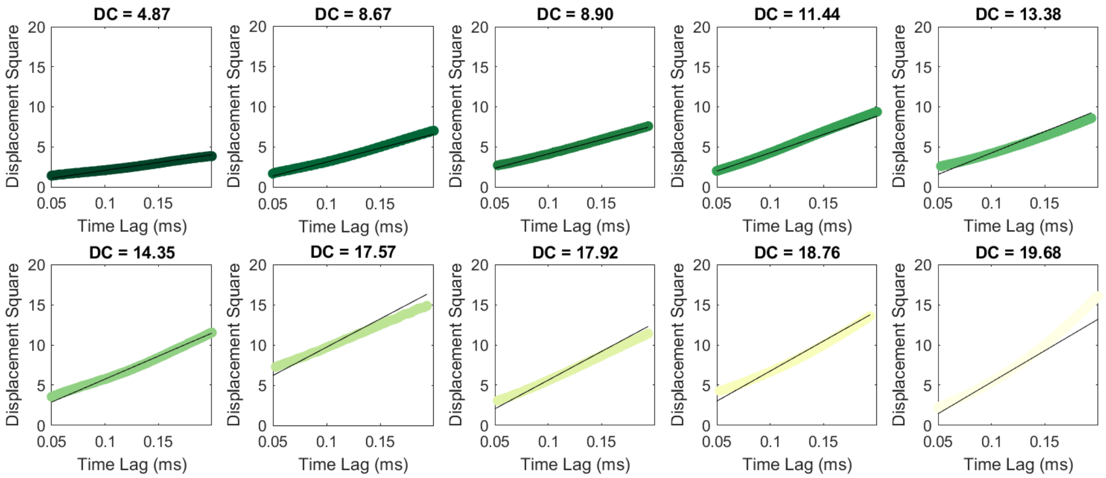


**Fig. S 10**: **Linear regression fits of drift displacement square.** Individual subject's drift displacement square as a function of time lags. Regression fits are also shown and diffusion coefficients, calculated based on the regression fits, are reported on top. Each color represents a different subject, ordered from lowest to highest diffusion constant.
